# A case study on the detailed reproducibility of a human cell atlas project

**DOI:** 10.1101/467993

**Authors:** Kui Hua, Xuegong Zhang

## Abstract

Reproducibility is a defining feature of a scientific discovery. Reproducibility can be at different levels for different types of study. The purpose of the Human Cell Atlas (HCA) project is to build maps of molecular signatures of all human cell types and states to serve as references for future discoveries. Constructing such a complex reference atlas must involve the assembly and aggregation of data from multiple labs, probably generated with different technologies. It has much higher requirements on reproducibility than individual research projects. To add another layer of complexity, the bioinformatics procedures involved for single-cell data have high flexibility and diversity. There are many factors in the processing and analysis of single-cell RNA-seq data that can shape the final results in different ways. To study what levels of reproducibility can be reached in current practices, we conducted a detailed reproduction study for a well-documented recent publication on the atlas of human blood dendritic cells as an example to break down the bioinformatics steps and factors that are crucial for the reproducibility at different levels. We found that the major scientific discovery can be well reproduced after some efforts, but there are also some differences in some details that may cause uncertainty in the future reference. This study provides a detailed case observation on the on-going discussions of the type of standards the HCA community should take when releasing data and publications to guarantee the reproducibility and reliability of the future atlas.

## Introduction

The Human Genome Project (HGP) has provided a complete list of virtually all nucleic acid sequences of the human genome [1, 2]. Such a list, together with the annotations completed by HGP as well as follow-up projects like ENCODE, provided a fundamental reference for current biological and medical studies on human [3-8]. The reference is a blueprint program for all cells in a human body. Not all cells in the same human body implement the program in the same way, and it is crucial to understand the genomic, transcriptomic, epigenomic, proteomic and metabolomics characteristics of all different types of cells at different time and organs to build a completing understanding of the human body in health and diseases. The Human Cell Atlas (HCA) program initiated from 2016 is the worldwide collaborative effort toward this goal [9, 10]. The program was largely enabled by the recent development of single-cell genomics, especially single-cell RNA-sequencing (scRNA-seq) technology. It provides efficient ways to measure the expression of thousands of genes in each of thousands to millions single cells [11]. Other single-cell technologies like single-cell ATAT-seq, single-cell Hi-C, single-cell metabolomics and technologies for single-cell resolution spatial transcriptomics are also under rapid development [12-16]. These single-cell technologies provide detailed observations on cells from multiple angles. They convert a biological cell to a data point in the high-dimensional space of multiple omics vectors. Cells that belong the same cell type or share similar molecular signatures may occupy a closed or open region in the high-dimensional space as clusters or distributions, and cell developments and state transitions can be reflected as linear or nonlinear trajectories in the space or in a certain subspace.

As a large-scale fundamental scientific program, HCA shares many features with HGP. But HCA has many unique and more challenging characteristics. Unlike the HGP which as a very specific and well-defined goal from the very beginning, for HCA, specifying what should be the essential defining components of the cell atlas and how they should be represented is one of the key questions to be answered by the program. There is a quick accumulation of publications on single-cell genomics studies in different human tissues [17-21]. They brought many advancements in the study of our immune systems, neuron systems, etc and also in early embryonic development. Some of these data and discoveries are becoming the earliest parts of the future human cell atlas. The most typical form of these discoveries are novel cell types or subtypes that have physiological or pathological importance, and gene expression markers of the discovered cell types [20, 22, 23]. Usually the discoveries are described in the published papers and supplementary materials, and some authors also release their raw or processed gene expression matrix data. There are major ongoing efforts in the HCA community for building Data Coordination Platforms to serve as future banks for HCA data [24]. Multiple steps of bioinformatics processing and high-level analysis are involved in all single-cell studies. The bioinformatics methods are usually described in the supplementary materials of published papers as they do for wet-lab protocols, mostly for the readers to understand what they have done. Most readers read the method parts to find hints to help their work on in-house data, without strong interests in reproducing published results. Many factors in the bioinformatics can have dramatic influences in the analysis of single-cell data, but a comprehensive guideline for how such factors should be documented and controlled for HCA has not been available yet.

After several attempts to work on the data of some recent work in this field, we realized that reproducing the final results from the original data can often be a challenging task. This may not imply any flew in the published work, but usually the information provided is not sufficient for reproduction. To study the major factors in bioinformatics pipelines that can influence various results, and the level of disclosure that is necessary to guarantee full reproducibility, we chose a well-documented recent publication on the atlas of human blood dendritic cells [23] as an example to conduct a breakup reanalysis on the data. The level of details provided with this paper is among the most sufficient ones in recent publications in this field.

We found that with this level of details disclosed, the major scientific conclusion can be well reproduced, but still with noticeable differences in some potentially important details. This detailed case study provides experimental observations to support the design of guidelines and standards of the HCA community on the documentation and quality control of bioinformatics pipelines.

## Results

The original paper updated the taxonomy of human blood dendritic cells (DCs) and monocytes by finding new cell types in a single-cell study of ∼2400 cells. In brief, they identified a new subtype of DC (named DC5), a new subdivision of a known subtype of DC (named DC2 and DC3), the existence of a conventional dendritic cell (cDC) progenitor, and two additional subtypes of monocytes by collecting and analyzing scRNA-seq data of 768 DCs and 372 monocytes derived from a healthy individual. The existence of these cell types are further confirmed in a subsequent study of additionally cells. They also studied the function of these newly discovered cell types and revealed the relationship between some of them. We mainly focused on the new discoveries on the taxonomy of the DCs in this reproduction experiment, i.e., the discovery of DC5 and the separation of previously known CD1C+ to the new subtypes DC2 and DC3 (Figure 1A).

**Figure 1.**
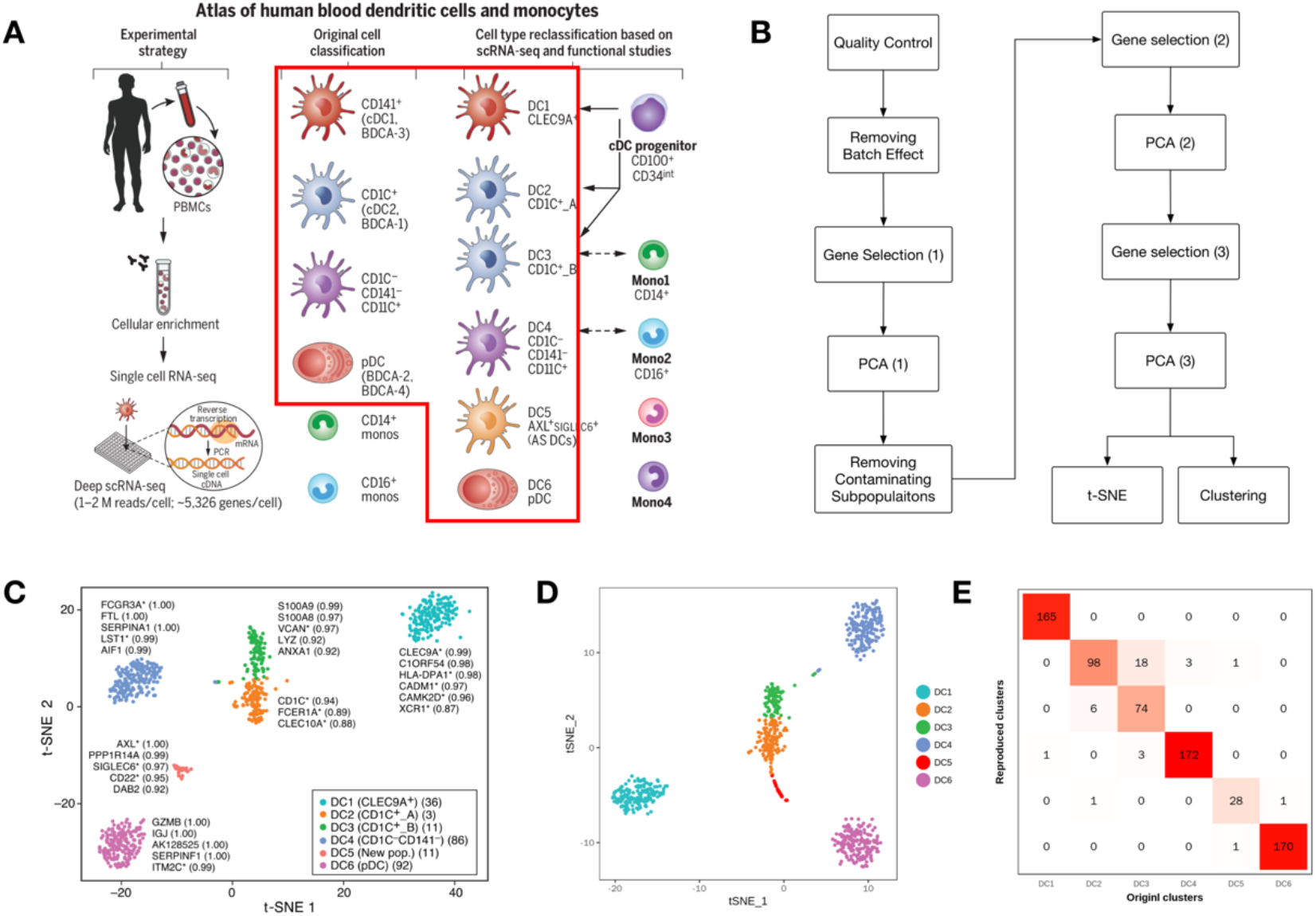
**(A)** Overview of the original work from its paper. The red frame shows the focus of this reproduction study. **(B)** Workflow of the computational part in the original work (extracted from the code given in the original supplementary materials). **(C)** T-SNE plot of DCs from the original paper. **(D)** Reproduced t-SNE plot of DCs. **(D)** Confusion matrix between the reproduced label of DCs and the label provided by the original paper.

The original paper had provided not only the expression data matrix, but also detailed information on the bioinformatics pipeline they used, including description of the analyzing procedures and the computer codes for the key analyzing steps. With all these materials, the reproduction procedure is quite straightforward, except for a few minor bugs that need to be fixed (see Supplementary Materials) [25]. Figure 1B shows a diagram of the bioinformatics pipeline we adopted in this experiment according to the description in the original paper and supplementary file.

Figures 1C–1E show the comparison of the t-SNE plot obtained in the reproduction experiment (Fig.1D) and the original one (Fig.1C). The major discovery that the DCs are of 6 clusters is well re-discovered. The two sets of clusters can be well aligned except some confusion between clusters DC2 and DC3, and a few tiny confusions between several other clusters. The relative relations of the 6 clusters are also mostly recovered, except that the new cluster DC5 is shown to have closer relations to the cluster DC2, which is not shown in the original t-SNE map. (Note that the t-SNE map does not preserve global distance information so differences in the overall layout of the plot do not indicate inconsistency [26-28].)

The major noticeable difference between the reproduced result and the original one is the separation of DC2 and DC3. Figure 2A and 2C show the comparison of the two heatmaps, drawn with the selected marker genes of each cluster with discriminating power of AUC>=0.85. The reproduction experiment assigned more cells to DC2 and less cells to DC3. From the enlarged heatmap (Figure 2B), we can see that the boundary between DC2 and DC3 on the marker genes are quite blurred. The slight differences in the clustering boundaries affected the identification of marker genes (There can also be other factors that caused the difference, see Supplementary Materials). To see how differences in cluster assignments would affect downstream analyses, we conducted standard differential expression genes (DE gene) analysis. Identifying DE genes is an old but still challenging question in single cell studies. Different methods can give quite different results [29,30]. Here we adopted the ‘one *vs* the rest’ strategy and selected DE genes based the AUC score as in the original paper. The table in Figure 2D summarizes the numbers of marker genes discovered for each cluster in the original paper and in the reproduction experiment. We can see although most of the marker genes ranked on the top are shared in the two experiments, the two gene lists do have some differences. The list of marker genes and their expression distributions in the two studies are shown in the Supplementary Materials.

**Figure 2.**
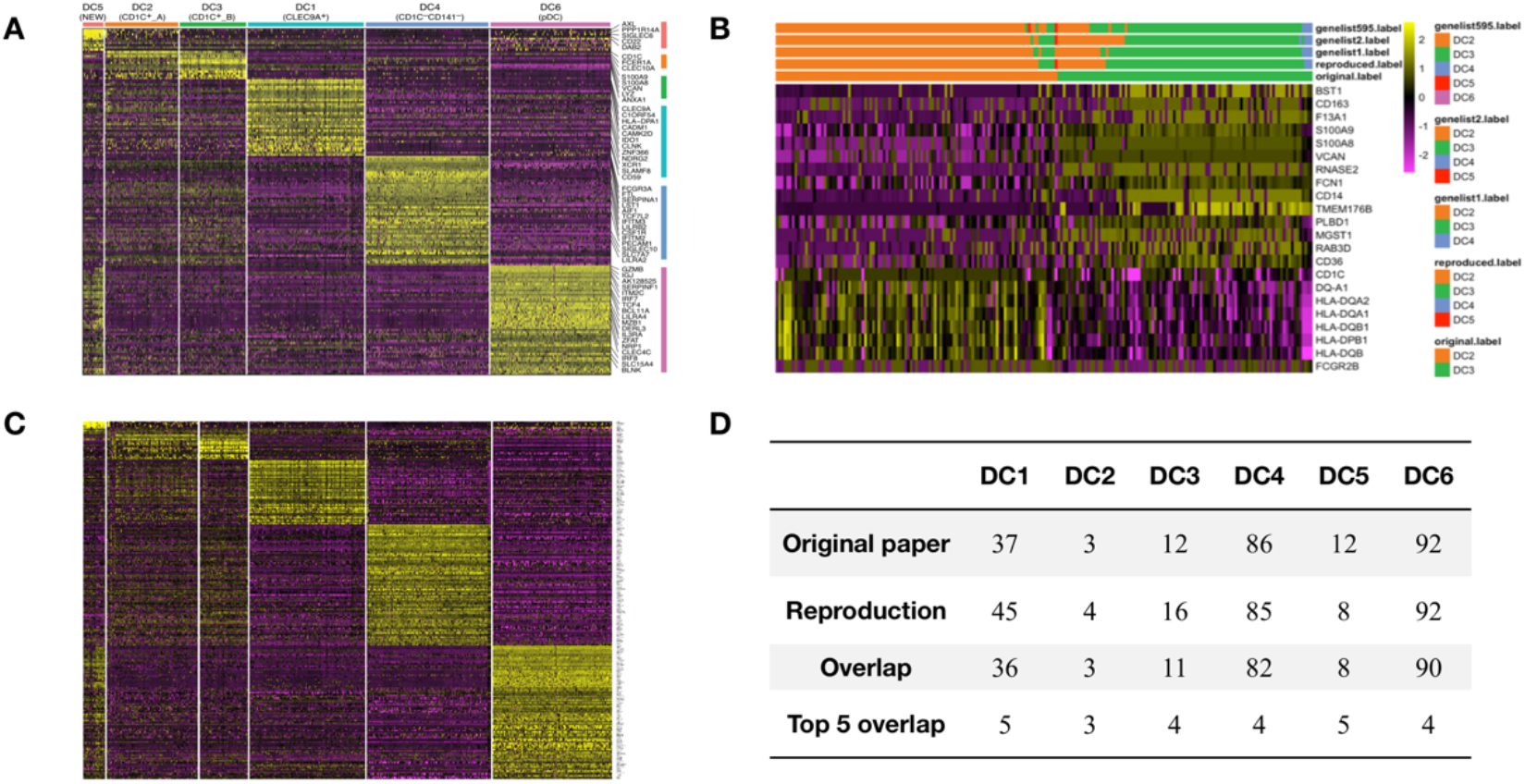
**(A)** Heatmap of cluster-specific markers from the original paper. **(B)** Heatmap of some markers of DC2 and DC3 (markers are selected according to original fig.2A). Side color bars illustrate clustering results using different gene lists. **(C)** Reproduced heatmap of cluster-specific markers. **(D)** Comparison of cluster-specific markers between the original results and reproduced results.

We conducted a step-by-step comparison of the reproduction experiment with the original experiment to pinpoint the factors that can cause the observed differences. The detailed description and analysis of these steps are provided in the Supplementary Materials.

The original paper did several downstream experiments with the discovered 6 DC clusters. They can be seen as examples for future applications of the 6 DC clusters as the reference of human DC atlas. In Figure 3A (the original Figure 3B), they showed the relations of the 4 monocyte clusters with the 6 DC clusters. Figure 3B shows the same analysis in the reproduction study. The relationship among mono1, mono2, mono3 and mono4 are basically preserved, but it is interesting to notice that with the presence of the monocyte cells, the DC5 cluster in the original study became connected with DC2, but it becomes separated from DC2 in the reproduction.

**Figure 3.**
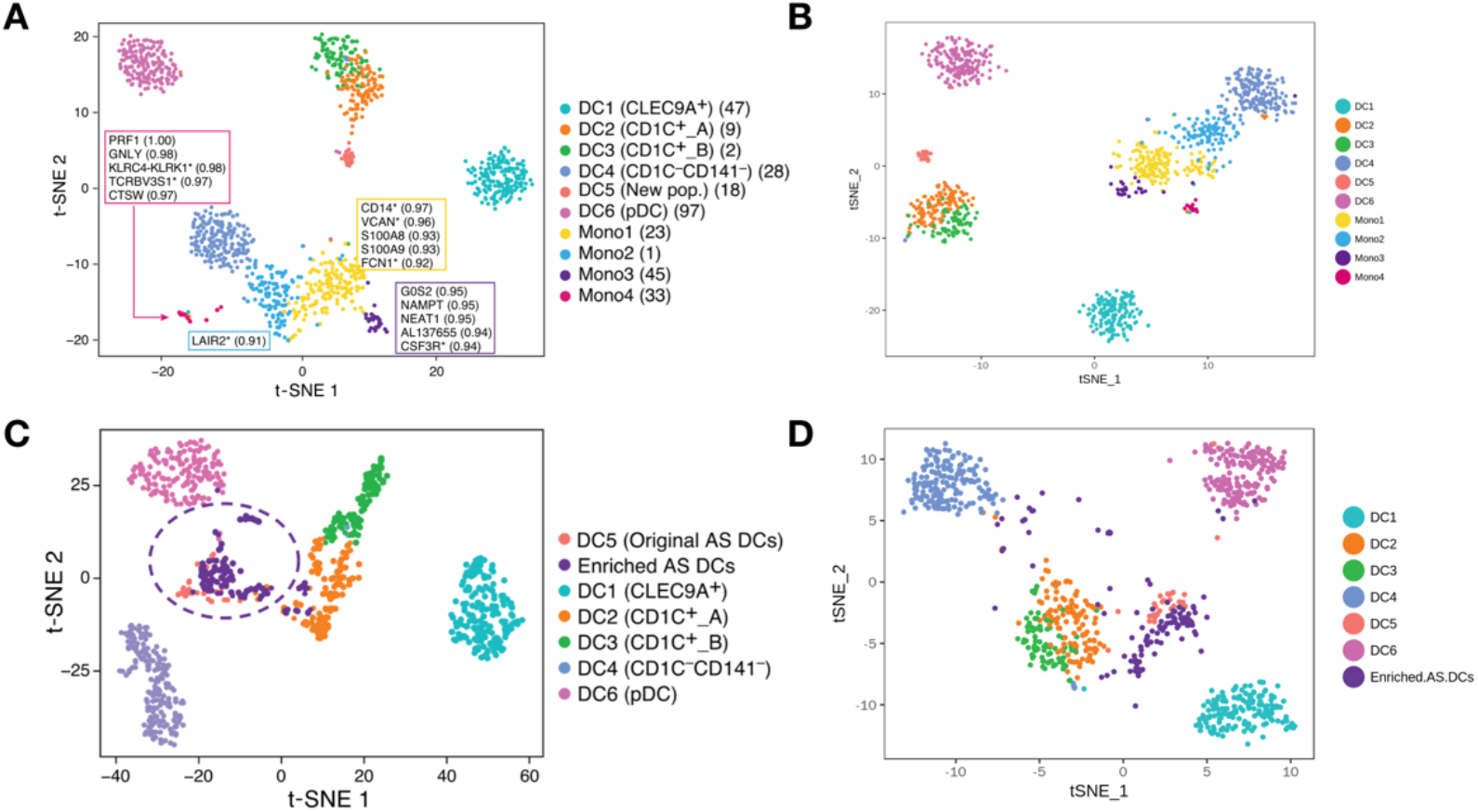
**(A)** Original figure 3B. **(B)** Reproduction of the original figure 3B. **(C)** Original figure 4C. **(B)** Reproduction of the original figure 4C. **(D)** Original figure 6G.

The original paper recruited 10 independent donors to confirm the existence of AXL +SIGLEC6+ cells (“AS DCs”), and used Figure 3C (original Fig.4C) to show their relation with the 6 DC clusters. Figure 3D shows the reproduction result. We can see the although the two t-SNE maps seem not identical, but the relationships revealed between the clusters are consistent, including the observed closeness of the AS DCs to the DC5 cluster.

**Figure 4.**
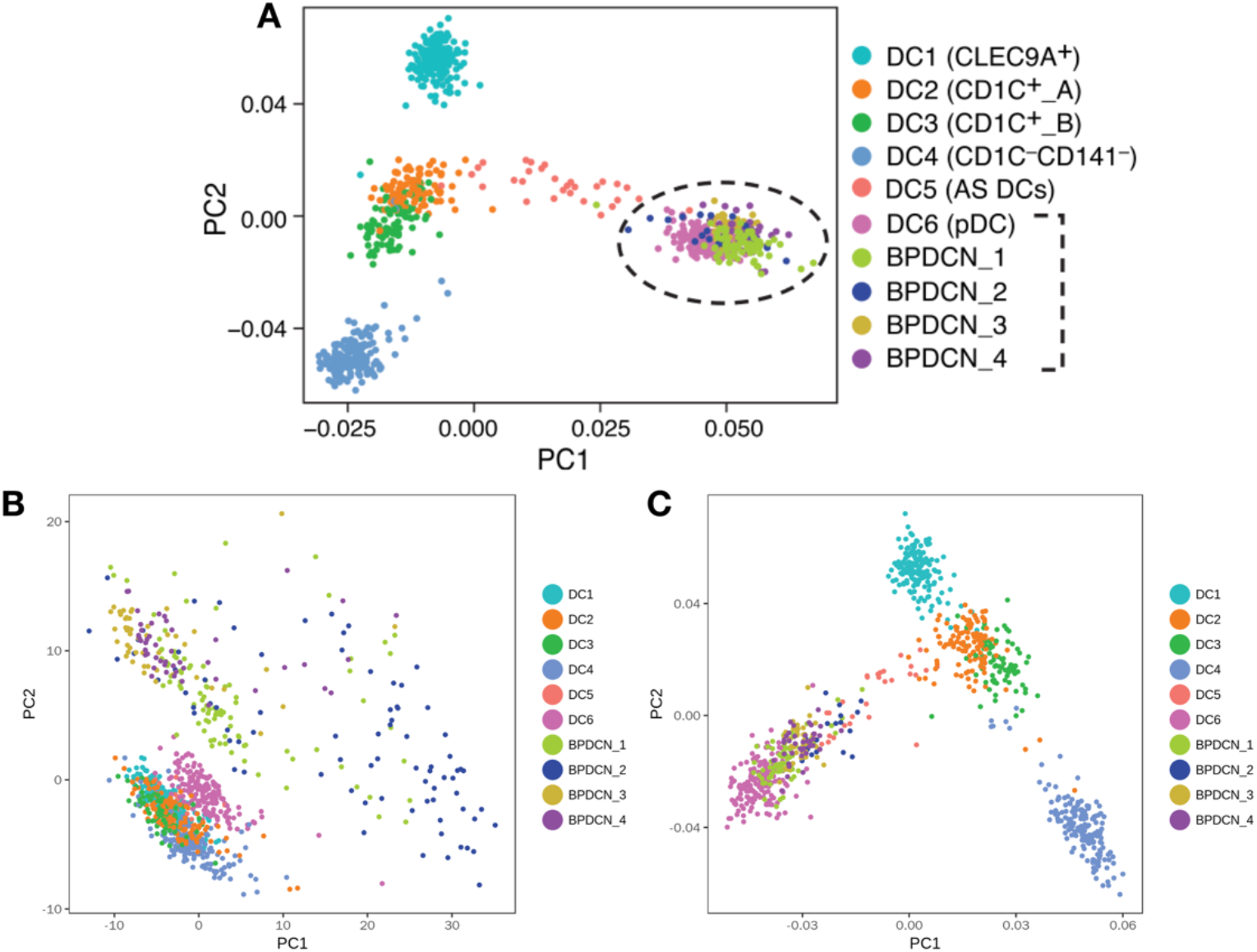
**(A)** Original figure 6G. **(B)** Reproduction of the original figure 6G using all BPDCN cells (n=269). **(C)** Reproduction of the original figure 6G using filtered BPDCN cells (n=174).

The original paper mapped pathogenic cells from blastic plasmacytoid dendritic cell neoplasm (BPDCN) patients to the healthy DC atlas. Figure 4A (original Fig.6G) showed that the BPDCN cells are mixed with DC6 cells in the PCA map. This result was generated using 174 out of 269 BPDCN cells. Since neither the final list of the used cells nor the filtering details were provided, we first used all 269 BPDCN cells to do the reproduction. In Figure 4B, we can see that cells from the 4 BPDCN patients are scattered aside the DCs in the map. They are close to DC6 but not mixed with it. We observed that the overall layout of the BPDCN cells are quite scattered, which also causes the 6 DC clusters squeezed, but some of the BPDCN cells seem to form a denser cluster which tend to be closer to DC6 than the more scattered ones. We therefore removed those scattered BPDCN cells as suspected low quality cells and redid the PCA (see supplementary for details). After this removing step, we got a similar PCA map where BPDCN cells are mixed with DC6 as in the original result. We noted that the description on the bioinformatics processing was less detailed for this part of the experiment than for the data used in the major discovery part in the original paper. Although we managed to get a similar result, this reproduction was achieved mostly by guessing and retries.

## Discussions

Single-cell genomics studies involve multiple steps of bioinformatics processing, including filtering of cells and selection of genes for quality control, batch-effect correction, normalization and dimension reductions for extracting major information, clustering, visualization and marker selection, etc. Some of the steps may need to be ran more than once and need to be adapted according to the data and the specific question. Due to the nature of variability between many biological problems and between single-cell genomics technologies, it may not be feasible to fix a “standard” bioinformatics pipeline. Therefore, providing standard descriptions of the pipeline with sufficient details is crucial for the reproducibility. Through this detailed reproduction case study, we observed that the extraordinarily detailed information of the bioinformatics procedures provided in the selected work has guaranteed the reproduction of the major scientific discovery, but still there are some unneglectable differences in the reproduced results. Some of the differences can be due to version changes of some third-party packages, and some may still be due to insufficient record or description in some minor details. Some seemingly subtle inaccuracies in the early and intermediate steps of processing may cause noticeable differences in the downstream results.

The HCA program is just at its beginning phase. All the current exciting discoveries based on single-cell genomics are just the beginning of the footstone construction of the future skyscraper. This distinguishes HCA projects with other single-cell studies that aim for only answering specific scientific questions. The construction of the full atlas will be done by joint efforts of many labs throughout the world, and it may span years or even decades. New discoveries are the focus of each individual publication, but the data used to support the discoveries are what accumulate in HCA, which will become part of the reference atlas for all future studies. The level of differences between results we observed in this case study can be subtle in terms of the major scientific discovery, but such tiny differences will be enlarged when the data act as part of the earliest footstones in the construction. There have been many recent discussions on conventions that are needed for journal publications to guarantee the reproducibility of scientific discoveries [31-33]. From the observations of this case study, it is evident that an even higher standard of reproducibility is needed for HCA as a reference for future sciences.

The importance of HCA is comparable with HGP but the scale is much larger. At the beginning of HGP, scientists had estimations of the final goal and had estimations on how much are still remaining during the progress of the program. For HCA, the final goal is more open and is of multifaceted nature. And the technology is evolving and expanding in revolutionary paces. For many studies, it is too early to judge which part of the data or intermediate results are unimportant details. At this phase, keeping a reasonable level of diversity of bioinformatics pipelines and methods is healthy for the community. But this should be accompanied by sufficient record and sharing of the details to guarantee full reproduction. In this reproduction experiment, to guarantee all results we got can be reproduced again by a third-party, we have created a notebook recording all details of the bioinformatics processing together with the corresponding live codes. Such efforts for guaranteeing reproducibility can serve as examples for all HCA projects. We propose building more stringent guidelines and standards on the information that need to be provided along with publications for projects that will be taken as part of the HCA program. And a mechanism of systematic third-party reproduction validation should be introduced.

## Data Availability

All results and the major bioinformatics processing are described in the Supplementary File. Data used in this reproduction and a jupyter notebook with all details of the reproduction are available at https://github.com/XuegongLab/HCA-reproducibility.

## Acknowledgement

We thank Nir Hacohen, Alexandra-Chloé Villani and Orit Rozenblatt-Rosen (authors of the original paper) for their helpful discussion. This work is supported by CZI Human Cell Atlas Pilot Project and the National Natural Science Foundation of China (61673231 and 61721003).

## References

[1] J. D. Watson, “The human genome project: past, present, and future,” Science, vol. 248, no. 4951, pp. 44–9, Apr 6 1990.

[2] F. S. Collins, M. Morgan, and A. Patrinos, “The Human Genome Project: lessons from large-scale biology,” Science, vol. 300, no. 5617, pp. 286–90, Apr 11 2003.

[3] C. International HapMap, “The International HapMap Project,” Nature, vol. 426, no. 6968, pp. 789–96, Dec 18 2003.

[4] E. P. Consortium, “The ENCODE (ENCyclopedia Of DNA Elements) Project,” Science, vol. 306, no. 5696, pp. 636–40, Oct 22 2004.

[5] J. L. Haines et al., “Complement factor H variant increases the risk of age-related macular degeneration,” Science, vol. 308, no. 5720, pp. 419–21, Apr 15 2005.

[6] C. Genomes Project et al., “A map of human genome variation from population-scale sequencing,” Nature, vol. 467, no. 7319, pp. 1061–73, Oct 28 2010.

[7] C. Genomes Project et al., “An integrated map of genetic variation from 1,092 human genomes,” Nature, vol. 491, no. 7422, pp. 56–65, Nov 1 2012.

[8] M. Kellis et al., “Defining functional DNA elements in the human genome,” Proc Natl Acad Sci U S A, vol. 111, no. 17, pp. 6131–8, Apr 29 2014.

[9] A. Regev et al., “The Human Cell Atlas,” (in English), Elife, vol. 6, Dec 5 2017.

[10] O. Rozenblatt-Rosen, M. J. T. Stubbington, A. Regev, and S. A. Teichmann, “The Human Cell Atlas: from vision to reality,” Nature, vol. 550, no. 7677, pp. 451–453, Oct 18 2017.

[11] V. Svensson, R. Vento-Tormo, and S. A. Teichmann, “Exponential scaling of single-cell RNA-seq in the past decade,” Nat Protoc, vol. 13, no. 4, pp. 599–604, Apr 2018.

[12] D. A. Cusanovich et al., “Multiplex single cell profiling of chromatin accessibility by combinatorial cellular indexing,” Science, vol. 348, no. 6237, pp. 910–4, May 22 2015.

[13] T. Nagano et al., “Single-cell Hi-C reveals cell-to-cell variability in chromosome structure,” Nature, vol. 502, no. 7469, pp. 59–64, Oct 3 2013.

[14] R. Zenobi, “Single-cell metabolomics: analytical and biological perspectives,” Science, vol. 342, no. 6163, p. 1243259, Dec 6 2013.

[15] N. Crosetto, M. Bienko, and A. van Oudenaarden, “Spatially resolved transcriptomics and beyond,” Nature Reviews Genetics, vol. 16, no. 1, pp. 57–66, Jan 2015.

[16] E. Lein, L. E. Borm, and S. Linnarsson, “The promise of spatial transcriptomics for neuroscience in the era of molecular cell typing,” Science, vol. 358, no. 6359, pp. 64–69, Oct 6 2017.

[17] S. Zhong et al., “A single-cell RNA-seq survey of the developmental landscape of the human prefrontal cortex,” Nature, vol. 555, no. 7697, pp. 524–528, Mar 22 2018.

[18] M. J. Muraro et al., “A Single-Cell Transcriptome Atlas of the Human Pancreas,” Cell Syst, vol. 3, no. 4, pp. 385–394 e3, Oct 26 2016.

[19] S. Darmanis et al., “A survey of human brain transcriptome diversity at the single cell level,” Proc Natl Acad Sci U S A, vol. 112, no. 23, pp. 7285–90, Jun 9 2015.

[20] E. Z. Macosko et al., “Highly Parallel Genome-wide Expression Profiling of Individual Cells Using Nanoliter Droplets,” Cell, vol. 161, no. 5, pp. 1202–1214, May 21 2015.

[21] M. J. T. Stubbington, O. Rozenblatt-Rosen, A. Regev, and S. A. Teichmann, “Single-cell transcriptomics to explore the immune system in health and disease,” Science, vol. 358, no. 6359, pp. 58–63, Oct 6 2017.

[22] D. A. Jaitin et al., “Massively parallel single-cell RNA-seq for marker-free decomposition of tissues into cell types,” Science, vol. 343, no. 6172, pp. 776–9, Feb 14 2014.

[23] A. C. Villani et al., “Single-cell RNA-seq reveals new types of human blood dendritic cells, monocytes, and progenitors,” Science, vol. 356, no. 6335, Apr 21 2017.

[24] H. D. team, “Data Coordination - Human Cell Atlas,” (https://www.humancellatlas.org/data-sharing).

[25] A. Butler, P. Hoffman, P. Smibert, E. Papalexi, and R. Satija, “Integrating single-cell transcriptomic data across different conditions, technologies, and species,” Nat Biotechnol, vol. 36, no. 5, pp. 411–420, Jun 2018.

[26] L. van der Maaten and G. Hinton, “Visualizing Data using t-SNE,” Journal of Machine Learning Research, vol. 9, pp. 2579–2605, Nov 2008.

[27] F. V. Martin Wattenberg, and Ian Johnson, “How to use t-sne effectively,” Distill, 2016.

[28] T. O. S. Yuansheng Zhou, “Using global t-SNE to preserve inter-cluster data structure,” bioRxiv, 2018.

[29] Miao, Zhun, and Xuegong Zhang. “Differential expression analyses for single-cell RNA-Seq: old questions on new data.” Quantitative Biology 4, no. 4 (2016): 243–260.

[30] Miao, Zhun, Ke Deng, Xiaowo Wang, and Xuegong Zhang. “DEsingle for detecting three types of differential expression in single-cell RNA-seq data.” Bioinformatics 1 (2018): 2.

[31] M. Baker, “IS THERE A REPRODUCIBILITY CRISIS?,” Nature, 2016.

[32] J. Berg, “Progress on reproducibility,” Science, 2018.

[33] P. B. Stark, “No reproducibility without preproducibility,” Nature, 2018.

